# Additivity of the genetic load is a product of both synergistic and antagonistic epistasis in *Drosophila*

**DOI:** 10.1101/2025.11.04.686015

**Authors:** Imran Sayyed, Halle McNamara, Troy Day, Adam K. Chippindale

## Abstract

Epistasis has been theoretically implicated in several major evolutionary processes, including the evolution of sex and speciation, but empirical evidence for its impact is still lacking. We tested for epistatic interactions using a full-sib inbreeding design in *Drosophila melanogaster*. We examined the effects of deleterious mutations on two sequential components of life-history: pupal productivity and pupa-to-adult viability. We observed an accelerating decline in pupal productivity with increasing inbreeding coefficient, consistent with synergistic epistasis, whereas pupa-to-adult viability exhibited a decelerating decline, consistent with antagonistic epistasis. This decoupling led to an overall linear decline in the production of adult offspring with increasing inbreeding. Our findings suggest that epistatic interactions can be highly trait-specific, underscoring the importance of analyzing individual, physiologically interconnected, fitness traits to understand the role of epistasis in evolutionary processes.

## Introduction

Epistasis, defined as a non-additive interaction between two or more genes, is hypothesized to play a significant role in regulating long-range linkage disequilibrium, thereby shaping pleiotropic associations and the organization of conserved genes (Singhal et al., 2023). Such genetic mechanisms are purported to shape the evolution of several fundamental biological processes, including speciation, sexual reproduction and perhaps, aging, suggesting epistasis may be a major driving force in these phenomena (Charlesworth, 1998; de Visser & Elena, 2007; Kondrashov, 1988; Wachter et al., 2013; Wagner et al., 1994; Weinreich et al., 2005; Zhang et al., 2024). Epistatic interactions can be categorized as either synergistic or antagonistic. Synergistic epistasis occurs when the combined deleterious effect of interacting alleles on fitness is greater than the sum of their individual effects, relative to an additive or multiplicative model. Conversely, antagonistic epistasis describes a scenario where the combined deleterious effect is less than that predicted from the individual effects (Weinreich et al., 2005; Zhang et al., 2024). Such non-additivity can be observed relative to the additive model of fitness or multiplicative model when fitness is analyzed on a logarithmic scale. However, empirical investigations of epistasis have proven challenging and often produce equivocal results, as explained below.

Two approaches have been employed to investigate patterns of fitness decline potentially associated with epistatic interactions among deleterious mutations. The first approach involves mutation accumulation (MA) experiments, which allow for the investigation of fitness decline patterns due to the increasing effects of spontaneously arising deleterious mutations, either under relaxed selection or using mutagens (Dickinson, 2008; Elena & Lenski, 1997; Kondrashov, 1994). These newly accumulated mutations may interact, causing deviations from additive fitness decline patterns. However, the evidence from mutation accumulation experiments regarding fitness decline potentially related to epistasis among deleterious alleles remains varied in quality and equivocal in outcomes (Rivero et al., 2003; Visser et al., 1997; West et al., 1998; Whitlock & Bourguet, 2000).

Another approach to studying fitness decline patterns involves inbreeding depression, which can be explained by either the partial dominance hypothesis or the overdominance hypothesis. Under the partial dominance hypothesis, fitness decline occurs due to the increased homozygosity of pre-existing recessive deleterious mutations maintained within a population. In contrast, the overdominance hypothesis posits that fitness reduction arises from a loss of heterozygote advantage. Given that heterozygote advantage is typically observed in a small percentage of loci in an equilibrium population, it is generally assumed that the majority of fitness decline due to inbreeding occurs under the partial dominance hypothesis (Charlesworth et al., 1991; Crow, 1999; Hedrick, 2012; Roff, 2002).

Studies of inbreeding depression have reported conflicting evidence, generally showing either no epistasis or antagonistic epistasis for fitness (Ávila et al., 2006; Rand et al., 2006; Sharp & Agrawal, 2016; Sohail et al., 2017; Szafraniec et al., 2003). Some of these discrepancies may be attributable to experimental biases, such as the repeated analysis of the same lines throughout the inbreeding process, sampling bias from the loss of lines due to lethal recessive deleterious mutations, and the effects of purging. However, a recent study by Domínguez-García et al., (2019) provided empirical evidence of synergistic epistasis for productivity using a full-sib inbred design in both laboratory and natural populations of *Drosophila melanogaster*. While Domínguez-García et al. (2019) addressed some of the extant challenges, certain experimental limitations persisted in their experiments, such as only analysing surviving lines which potentially creates a bias against synergistic epistasis among strong deleterious mutations. Another frequently overlooked factor in studies of inbreeding and the pattern of fitness decline is the direct influence of the inbreeding coefficient on fitness components separately. A critical assumption often made is that the accumulation of deleterious mutations, and consequently their homozygous expression, increases at a consistent linear rate across a variety of fitness components affected by these mutations. Under such an assumption, the inbreeding coefficient is expected to affect various life history traits equally leading to a linear decline in the overall fitness.

While a linear relationship between increasing homozygosity of recessive deleterious mutations and their phenotypic impact is often assumed across all fitness components, it is plausible that different life history traits may exhibit varying rates or non-linear patterns of decline in response to increasing inbreeding coefficients (Charlesworth, 1998; Falconer, 1996; Willis, 1993). Investigating these potential differential responses in fitness components is crucial for a comprehensive understanding of genetic load and epistatic interactions.

This conflicted but developing literature inspired our study, in which we investigated the effect of increasing inbreeding coefficient on several composite traits which determine the fitness of fruit flies. We used a full-sib inbreeding design but replicated parental lines to create ten copies of each initiating genotype. We investigated the patterns of decline in pupal productivity, pupa-to-adult viability, and overall offspring productivity with increasing inbreeding coefficients. Our results revealed an accelerated decline in pupal productivity but a decelerated decline in pupa-to-adult viability. This culminated in an overall linear decline in offspring productivity with increasing inbreeding coefficients. These findings highlight the complexity of studying epistasis among deleterious mutations affecting fitness, especially given that fitness is a highly composite trait. Furthermore, our results underscore the fact that distinct epistatic interactions can influence specific life-history traits without necessarily altering the expected additive effects on overall fitness.

## Materials and Methods

### Study system

Our experimental design utilized a laboratory population of *Drosophila melanogaster* designated *Control* (C_1_) that is descended from the *Ives* (IV) base population. IV was established from a wild caught sample of 400 flies (1:1 sex ratio) in Amherst, Massachusetts in 1975 (Rose & Charlesworth, 1981a, 1981b). The C_1_ population has a varied demographic history but is outbred and had been maintained under standardized environmental conditions since 1980. These conditions include discrete generations, a population size of approximately 2000 individuals per generation, cultivation on a banana-agar-killed yeast food medium, a moderate larval density (80-100 larvae per vial), a constant temperature of 25°C, and a 12-hour light:12-hour dark photoperiod. This population had been maintained for over 150 generations on a two-week life cycle. With more than 400 total generations of controlled laboratory maintenance, the C_1_ population is expected to be at, or close to, an evolutionary equilibrium for standing genetic variation. This characteristic makes it particularly well-suited for estimating mutational load at an equilibrium and measuring fitness under conditions of evolutionary history by assessing various life-history traits.

### Full-sib Inbreeding

To establish the inbred lines, 30 unmated male and female pairs were isolated from the C_1_ population, with each pair initiating a distinct line within a separate vial. In the second generation, 10 unmated male and female pairs were isolated from the offspring of each established inbred line to create 10 replicate ‘sub-lines’ (300 lines in total). For the subsequent five generations, each sub-line was maintained by randomly collecting a single pair of unmated male and female offspring; a sub-line was terminated if no viable adult pairs were produced in a generation. This ‘sub-line’ design helps us to account for the possible extinction of lines in the analysis due to strong deleterious effect arising from synergistic epistasis leading to a bias against accelerated decline in surviving lines. In total, these experimental lines underwent six generations of inbreeding, during which fitness decline was assessed.

### Assessing fitness

The pattern of fitness decline due to increasing inbreeding was assessed by measuring several key life-history traits. These included pupal productivity as the number of pupae produced by an inbred pair, the pupae to adult viability during pupal stage, and offspring productivity measured as the total number of adults eclosed for each surviving sub-line. Concurrently, these same traits were measured in the outbred control population (C_1_) by sampling 50 male and female pairs each generation, synchronized with the measurements from the inbred lines. For any sub-line that did not survive in a given generation, the zero values were still treated as data in analyses.

### Analysis

All analyses were performed using R version 4.4.2 (R Core Team, 2024).

Since pupal productivity and offspring productivity are composite traits that account for the reproductive fitness of parents, number of eggs laid, hatched and pupated successfully, the expected inbreeding coefficient (*ICf*) for pupal productivity and offspring productivity was calculated and assigned for each generation of inbreeding by averaging the inbreeding coefficients of the mother and the offspring based on the previous work by Domínguez-García et al. (2019). This resulted in expected *ICf* values of 0, 0.125, 0.313, 0.438, 0.547, 0.633 for the generations 1 to 6, respectively, for the entire genome in a full-sib line. For pupa-to-adult viability and sub-line survival, the expected inbreeding coefficient was calculated based solely on the offspring’s inbreeding coefficient (*IC*), yielding values of 0, 0.25, 0.375, 0.5, 0.594, 0.671. Using the lme4 package in R, we fitted linear and quadratic hurdle models to the measured traits against the expected inbreeding coefficient. In our regression models, inbred lines were included as a random effect to account for stochasticity inherent to each homozygous line. Model comparisons between linear and quadratic fits for each trait were performed using Akaike Information Criterion corrected for small sample sizes (AICc), Bayesian Information Criterion (BIC), and log-likelihood comparisons to determine whether the observed decline was linear or non-linear with increasing inbreeding coefficient.

## Results

1. Survival of sublines: The number of surviving sub-lines per inbred line exhibited an accelerating decline of 58% over 6 generations with increasing inbreeding coefficient (*ICf*), as inferred from comparisons between quadratic and linear model fits (Figure 1, Table 1). This non-linear decline in sub-line survival suggests a significant effect of synergistic epistasis among lethal mutations or synergistic interactions between recessive deleterious mutations affecting sterility or pre-copulatory mortality.

**Table 1.** AICc, BIC and log-likelihood values of both linear and quadratic fitted models for each trait measured. P-value from Chi-square test of log-likelihood fits of respective model comparisons suggest whether quadratic model is significantly different than linear model or not.

| *Trait* | *Model* | *AICc* | *BIC* | *Log-Lik* | *Pr(>Chisq)* |
| --- | --- | --- | --- | --- | --- |
| *No. of Sublines*<br>*Surviving* | Linear | 649.4751 | 658.8825 | -321.67 | **1.931e-05 \*\*\*** |
|  | **Quadratic** | **633.3126** | **645.8085** | **-312.54** |  |
| *Pupal Productivity* | Linear | 1084.042 | 1099.282 | -539.01 | **0.01361 \*** |
|  | **Quadratic** | **1079.968** | **1100.281** | **-535.97** |  |
| *Pupa-to-adult*<br>*Viability* | Linear | -691.9706 | -676.7331 | -783.28 | **4.339e-06 \*\*\*** |
|  | **Quadratic** | **-715.0756** | **-694.7656** | **-772.72** |  |
| *Adult Progeny*<br>*Eclosed* | Linear | 951.3498 | 966.5873 | -472.66 | 0.7903 |
|  | Quadratic | 953.2925 | 973.6025 | -472.63 |  |

**Figure 1.**
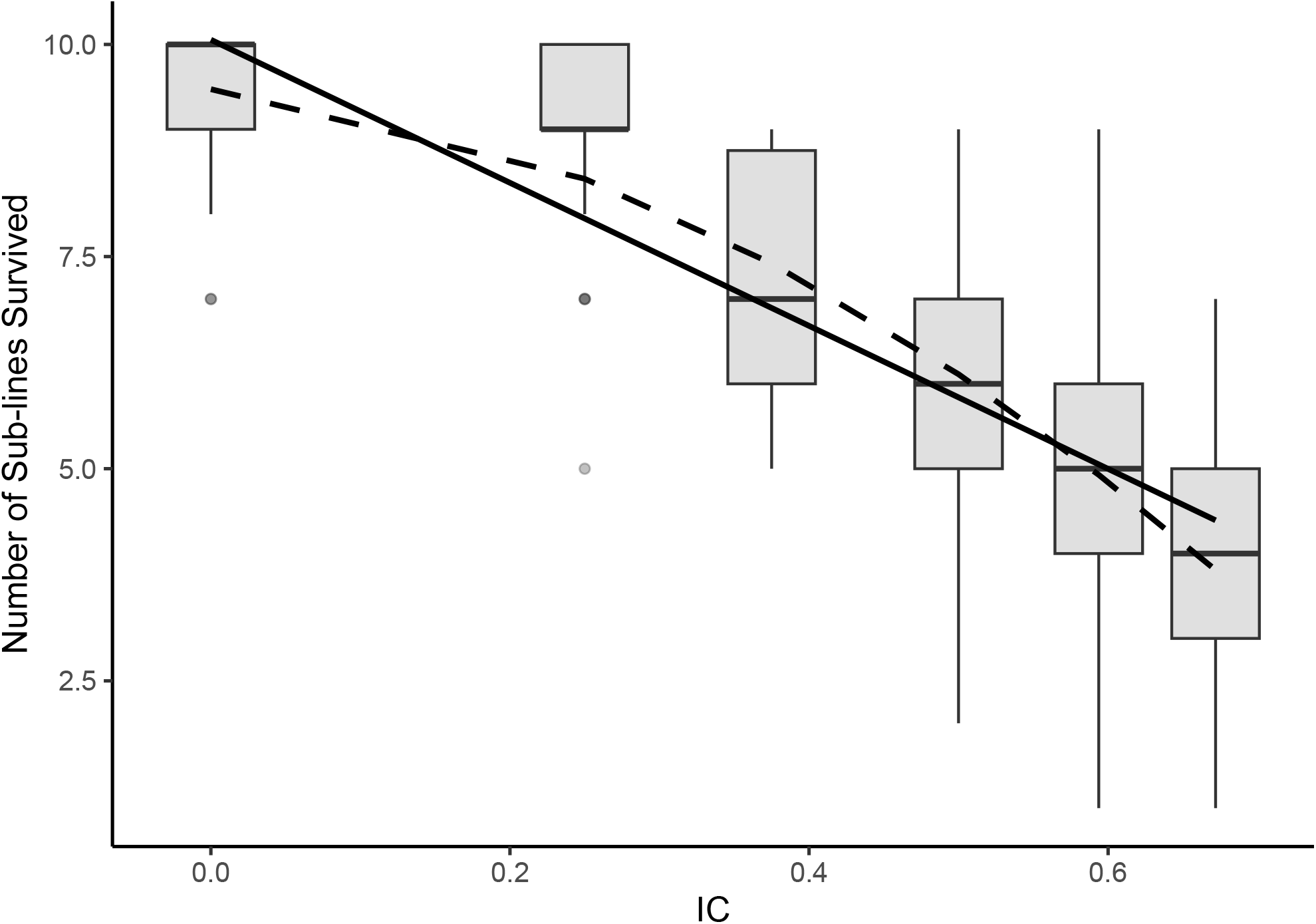
Boxplot of number of sublines surviving at a given IC. The black solid line represents the best linear fit, while dotted non-linear line represents quadratic fit for the decline in subline survival with IC.
2. Pupal productivity: Overall pupal productivity declined by 56.02% in a non-linear fashion with increasing inbreeding coefficient (Figure 2, Table 1). Among the 30 inbred lines, 24 lines showed a significant accelerated decline in pupal productivity, providing strong evidence for synergistic epistasis involving mutations that affect egg-to-pupae viability and development.

**Figure 2.**
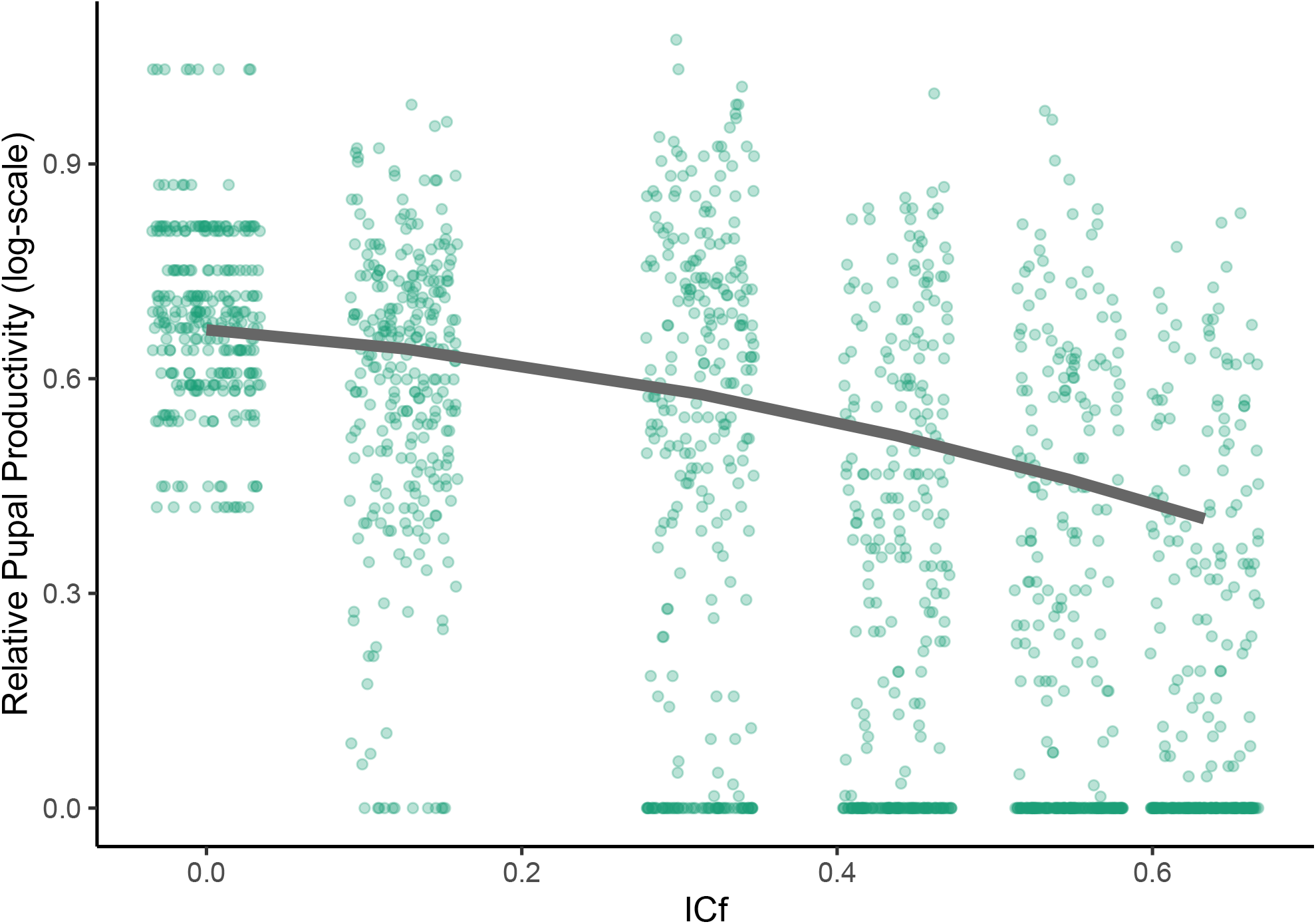
Quadratic regression for pupal productivity (on log-scale) with increasing average inbreeding coefficient (ICf). The relative scale is with respect to the outbred lines. Each point indicates pupal productivity for each inbred sub-line at a given ICf.
3. Pupa-to-adult viability: Overall pupa-to-adult viability, representing survival during the pupal stage of pre-adult development, declined by 18.65% in a non-linear fashion with increasing inbreeding coefficient (*IC*) (Figure 3, Table 1). Out of the 30 inbred lines, 22 lines exhibited a significant deceleration in pupa-to-adult viability, suggesting strong evidence for antagonistic epistasis among mutations affecting this developmental stage.

**Figure 3.**
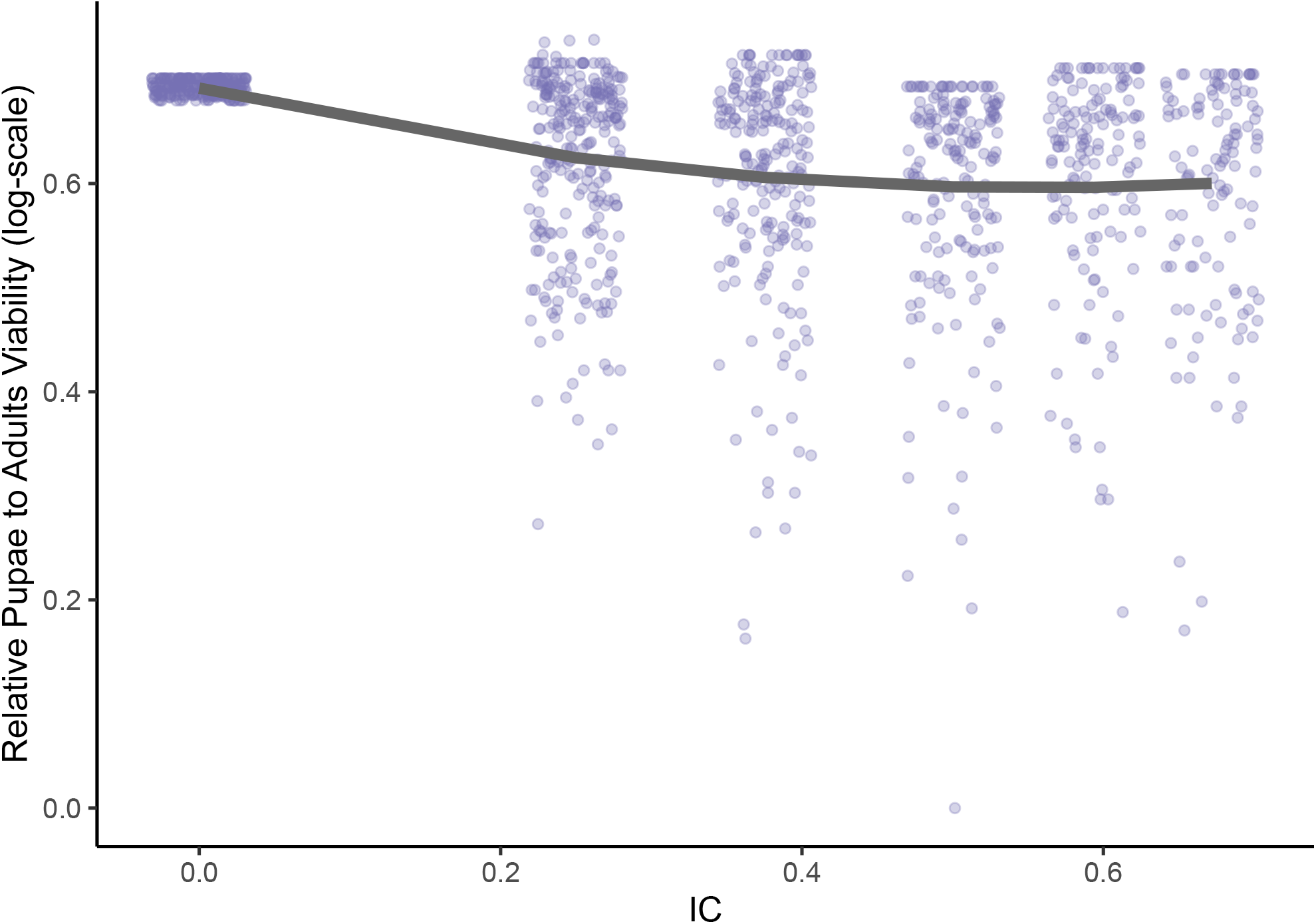
Quadratic regression for pupae-to-adult viability (on log-scale) with increasing inbreeding coefficient (IC). The relative scale is with respect to the outbred lines. Each point indicates pupae-to-adult viability for each surviving sub-line at a given IC, as this viability ratio can not be calculated for extinct lines.
4. Number of adults eclosed: The overall number of adults eclosing, representing survival through the entire pre-adult developmental stage, declined by 64.37% in a linear fashion with increasing inbreeding coefficient (Figure 4, Table 1). Among the 30 inbred lines, 27 lines showed a significant linear decline in the number of adults eclosed, which appears counter-intuitive given the results for pupal productivity and pupa-to-adult viability. Instead, this linear pattern can be interpreted as an outcome of the combined effects of accelerated decline in pupal productivity and de-accelerated decline in pupa-to-adult viability, resulting in an overall linear reduction in the number of adults eclosed with increasing *ICf*.

**Figure 4.**
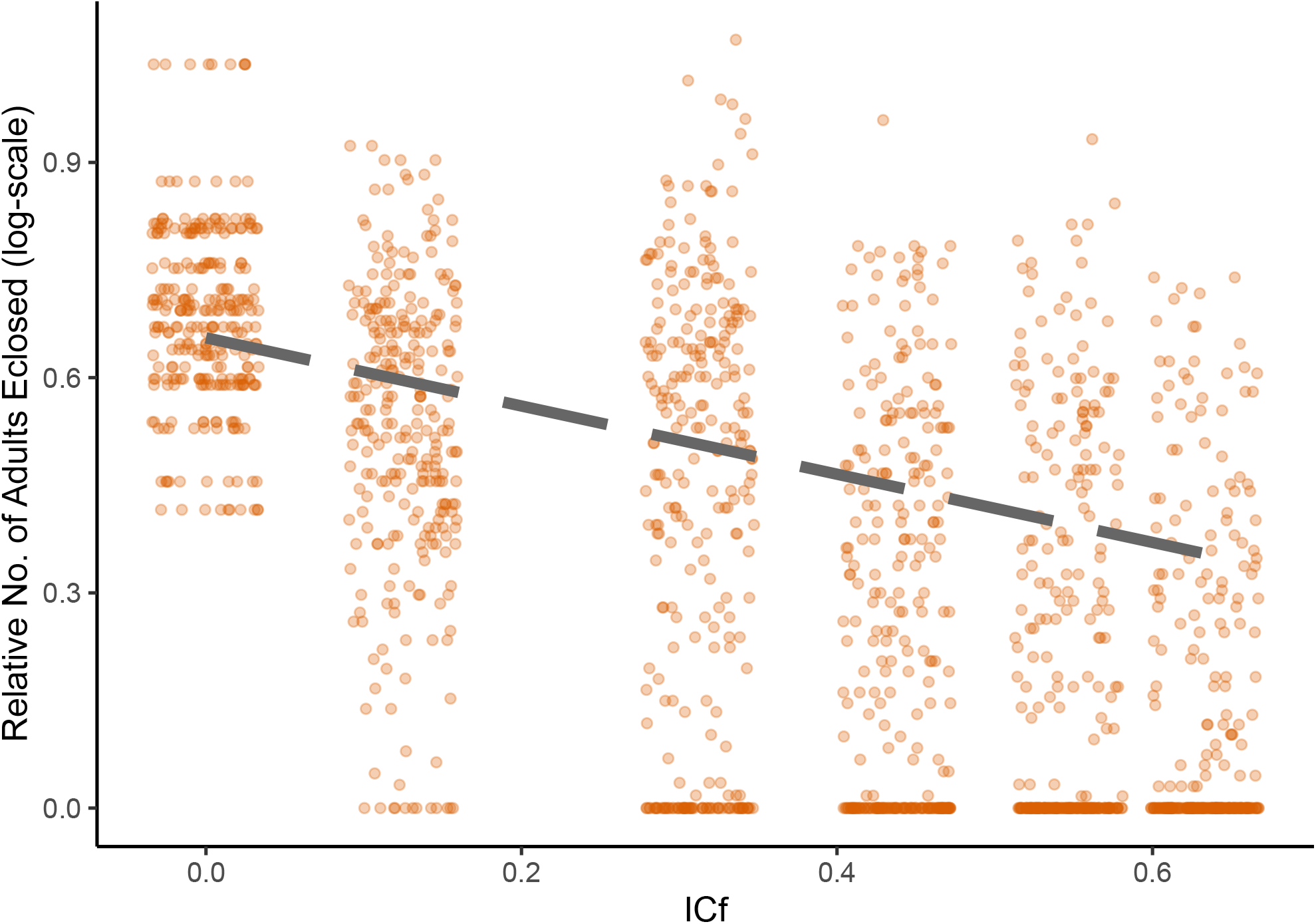
Linear regression for number of adults eclosed (on log-scale) with increasing average inbreeding coefficient (ICf). The relative scale is with respect to the outbred lines. Each point indicates number of adults eclosed for each inbred sub-line at a given ICf.

## Discussion

Experiments aimed at deducing the effects of increasing mutation load have yielded a range of outcomes and are equivocal, as a literature, on the question of epistasis for fitness effects. These studies have varied in scale and design, with limits to inference of any one approach. For example, mutation accumulation (MA) experiments have not provided consistent evidence for epistasis among deleterious mutations (Ávila et al., 2006; Dickinson, 2008; Elena & Lenski, 1997; Kibota & Lynch, 1996; Visser et al., 1997). These investigations typically focus on newly arising spontaneous mutations that would be subject to natural selection and subsequent purging from the population; such spontaneous mutations may not be the ones segregating naturally and having the greatest impact in populations. Consequently, an inbreeding approach offers a potentially valuable alternative approach for examining epistatic interactions among pre-existing recessive deleterious mutations that have accumulated in a population under mutation-selection balance in natural populations (Charlesworth & Hughes, 1996; Domínguez-García et al., 2019; Swindell & Bouzat, 2006). Our study population has been maintained for over 400 generations in the lab at large population size, a duration that is expected to result in something approaching an equilibrium state, minimizing the impact of purging and balancing selection on the analysis of epistatic interactions.

Few inbreeding studies have been directed at the question of epistasis, however several do suggest the presence of epistatic interactions, both synergistic and antagonistic, affecting various fitness-related traits (Domínguez-García et al., 2019; Rosa et al., 2005; Willis, 1993). Given that fitness is a complex, polygenic trait, and researchers often use proxies for total fitness, it is important to dissect epistatic interactions within its constituent traits individually, where feasible. Our current investigation was inspired by Domínguez-García et al. (2019) who suggested the existence of synergistic epistasis for fitness, evidenced by an accelerated decline in pupal productivity in *D. melanogaster* laboratory and natural populations. Consistent with their findings, our results also indicated an accelerating decline in a similar metric of pupal productivity. However, we also observed a deceleration in the effects of inbreeding on pupa-to-adult viability, which ultimately led to a linear decline in the total number of eclosed adults (total fitness) with increasing inbreeding.

Why then is the impact of inbreeding different for pupal productivity and pupal viability? Pupal productivity is a highly composite character, encompassing almost all aspects of total fitness: adult fertility in the parental generation and larval viability in the subsequent generation. Potentially the sheer complexity and number of different ways that productivity could fail through infertility or poor juvenile competition and development, with performance deficits in one phase potentially compounding in the next, creates the conditions for synergism across mutations. In contrast, pupal survival is likely to be an intensely canalized trait, with the metamorphic animal executing a highly choreographed developmental phase highly dependent on the interactions of rare alleles with environmental stress (Bendixsen et al., 2017; Borne et al., 2021; Collet et al., 2023).

In summary, although we are unable to pinpoint the driving elements from the current analysis, we speculate that the expression of deleterious mutations in one life stage could have a potential cascading or ‘synergistic’ effect in later stages leading to larger than expected fitness decline, without directly interacting with other deleterious mutations. This hypothesis involves condition-dependence, in which erosion of condition below thresholds results in catastrophic failures and morbidity. This mechanism may be complimentary to a study from Sohail et al. (2017), who detected rare loss-of-function alleles exhibiting synergistic epistasis under purifying selection in both human and *Drosophila* populations. On the other hand, it explains why purifying selection during mutation-selection balance could promote the accumulation of recessive deleterious mutations exhibiting antagonistic epistasis during the pupal stage in *D. melanogaster*. This hypothesis is consistent with evidence of a higher rate of non-synonymous substitutions relative to mutation rate (ω) in coding regions during the pupal stage of *D. melanogaster* compared to ω in the coding regions involved in pupal productivity (Coronado-Zamora et al., 2019).

Our findings demonstrate that epistatic interactions among deleterious mutations may be strongly trait-specific, to the point of shifting from synergistic to antagonistic depending upon the particular fitness components being assessed. This means that, at least in our study population, even measurements of fertility and survival that capture nearly all aspects of total fitness may misrepresent that parameter, which is of greatest importance from the theoretical standpoint. While synergistic epistasis for total fitness has been implicated in broadly important evolutionary phenomena, such as the evolution of sexual reproduction, our results do not support such a role. However, our study does raise intriguing questions about why different life stages and combinations of fitness characters would yield opposing results regarding epistasis. Hypotheses regarding the relationship between purifying selection, epistasis, and the ratio of non-synonymous to synonymous substitutions (ω) can serve as a starting point to investigate such trait-specific epistatic interactions, but their combination in sequence poses special challenges, particularly for highly composite traits. Overall, understanding the genetic and physiological underpinnings of non-additivity will allow a greater insight into the evolutionary architecture of fitness and into the genetic load of populations.

## Conflict of interest

The authors declare they have no conflict of interest.

## Data archiving

To be uploaded on https://datadryad.org/

## Acknowledgments

Funding for the work was from NSERC Discovery grants to AKC & TD. We thank all the undergraduate students involved in the population maintenance. We thank Harshavardhan Thygarajan, Mindy Baroody, Avery Want, Josh Kowal for their efforts in the lab.

## Author contribution statement

IS was responsible for designing the protocol, analysing data, creating figures and tables and interpreting results. IS and HM contributed equally to conducting the research and collecting the data. IS wrote the first draft of the paper. IS and AKC contributed to the updating of reference list and writing the report. IS, TD and AKC contributed to the final draft and revisions.

## Notes

### Competing Interest Statement

The authors have declared no competing interest.

